# Koalas, cameras and a primary school: Obtaining ecological insights for land management through a mini-citizen science project with young students

**DOI:** 10.1101/2023.06.22.546188

**Authors:** Sharon Williams, Gerard Salmon, Daniel Dempsey, David Philpot, Geoff Faulkner

**Affiliations:** Science specialist at Goodna State School; Goodna State School during citizen science project; Downer - website and server developer for project; Faulkner Consulting

**Keywords:** primary, students, citizen-science, koala, fauna, assessment

## Abstract

Koalas are an endangered species in Eastern Australia and the management of habitat revegetation sites is essential for the marsupial’s survival. This report summarises the results of a partnership between revegetation managers and a local primary school that connected children to real world scientific practice. Ten infrared motion sensor cameras were installed on a five-year-old revegetation offset site in south-east Queensland, Australia to capture movements of koalas and other fauna. Primary school students screened images from these cameras for animal presence in a mini-citizen science project through an easy-to-use website created specifically for this project. Aspects of these results were used to modify the land’s management by site supervisors. This report outlines processes involved in having Queensland primary school students participate in a mini web-based citizen science project, presents some of the ecological results obtained through the investigation and makes recommendations for future projects of this type.

## Introduction

Australian koalas (*Phascolarctos cinereus)* are classified as an endangered species in Queensland, New South Wales and the Australian Capital Territory (Department of Agriculture Fisheries and Forestry 2022), and are at high risk of extinction by 2050 in Eastern Australia (WWF International 2022, 2019). In fact, the large-scale bushfires, and associated habitat destruction, in Australia in early 2020 have put even more pressure on koala populations, making the assessment of the successfulness of koala habitat revegetation sites an essential part of preserving the marsupial. Pressure on koala numbers in south-east Queensland comes from numerous sources including traffic related deaths and the prevalence of disease and infertility caused by the obligate intracellular bacteria *Chlamydia (Chlamydia pecorum* and *Chlamydia pneumonia)* (Polkinghorne, Hanger and Timms 2013). Recent studies have shown *Chlamydia* infection alters fertility in female and male koalas (Hulse et al 2021). It is hoped that new vaccine technology for *Chlamydia* will halt the decline in breeding success for this marsupial (Quigley and Timms 2021). However, given that the reduction of native tracts of habitable land due to urban industrial and rural development poses the largest threat (Department of Environment and Science 2022) successful off set management is essential in south east Queensland.

Under Queensland law, when koala habitat is removed by industry to construct housing estates, roads or railways, replacement revegetated land must be created to offset the damage and protect the wildlife (Department of Environment and Science 2021). Analysing the successfulness of these habitats is traditionally characterised by assessing the tree health and non-juvenile koala habitat tree numbers (as outlined chapter 3 of Department of Environment and Science 2020). Koala population numbers have been estimated by gathering evidence of wildlife presence through methods such as the monitoring of scat and the use of dogs to follow scents (Cristescu et al. 2015), and the counting of individual animals using sampling methods such as line transects (Dique et al. 2003). However, the use of motion sensor cameras has had limited use in the Greater Ipswich region of Queensland for such purposes due to the initial expense of purchasing the equipment, the time required to change over data cards and the labour-intensive nature of analysing the images.

The use of citizen scientists in research has become popular in the last decade, particularly in ecology, as it harnesses the observational skills of numerous members of the public. According to Dickinson et al. (2012) ecological research can benefit from citizen science projects by expanding spatial research and supplementing localised existing programs, particularly those focussed on the biological studies of global climate change, phenology, landscape ecology, and rare and invasive species distribution and populations (Dickinson et al. 2012). Examples of Australian citizen science projects have included monitoring humpback whale migration (Bruce et al. 2014), the identification of frogs through smartphone sound recognition apps (Rowley et al. 2019), and studies of Tasmanian suburban garden habitats that showed the positive impact household gardening practices can have on bird conservation (Daniels and Kirkpatrick 2006). Citizen science projects for koalas have included projects that have run over the past two decades and recorded public sightings of koalas in south-east Queensland, that informed koala distribution data collected by other means (Dissanayake et al. 2019). However, the use of primary school students as citizen scientists in koala research is limited.

Landscape ecology practitioners are concerned with how the composition of the landscape effects the diversity of organisms, their populations, and how well these living things survive and reproduce within a given location (Dickinson et al. 2012). This research project focussed primarily on koala numbers (and other fauna) within the revegetation site in Ipswich. When motion sensing or time-lapse cameras are installed in a habitat, hundreds of thousands of images are captured within relatively short periods of time. The laborious task of scanning images for the presence of fauna can be efficiently conducted when members of the public perform the preliminary scans, allowing the science researchers to devote their attention to smaller subsets of images that contain species of interest. When citizen scientists conduct preliminary scans, data can be collected on the activity of other fauna within the region beyond the key species of interest. In this project, motion sensor cameras photographed kangaroos, feral foxes and feral pigs, and native bird life. However, the involvement of primary school students as citizen scientists requires very simple, user-friendly interfaces so such young children can participate easily in the data gathering and analysis.

It is often contended that the potential for promoting student participation in science, technology, engineering and mathematics (STEM) programs is enhanced when connections to real world projects are made. Indeed, in combining research with public education, citizen science can address broader societal impacts by engaging non-scientists, and even students, in authentic research experiences especially given that access to modern communication tools such as smart phones, the internet and computers (Dickinson et al. 2012) which are readily available in most schools.

Contrastingly, financing and managing large numbers of students on field trips or excursions to private locations or national parks can be time consuming and ecologically damaging and thus the pattern has been to take smaller groups of talented students to science enrichment projects and report their findings back to their peers. However, to allow the whole student population of the Ipswich primary school, Goodna State school, (approximately 700 students) to engage with a real-world project, an Advance Queensland Engaging Science Grant was sought and awarded to the school in 2019. This fostered the creation of a citizen science project that allowed for both a select group of students to physically attend a koala revegetation site, and for the rest of the school to act as online citizen scientists and assist in the analysing of photographs from the area at Mutdapilly, Ipswich. This report summarises the fauna identified by students, including koalas, considers the success of this research partnership and reflects upon the effectiveness of primary school students as citizen scientists whilst highlighting some development and implementation challenges within a public-school setting. This report also speculates on the utility of similar projects in future ecological research.

## Materials and Methods

Ten Browning Dark Ops, Pro XD Model Number: BTC-6PXD, infrared, colour and motion sensor capable cameras were installed in key locations around the Mutdapilly offset site in south-east Queensland. The land was formerly cattle and dairy grazing country and was purchased for rehabilitation into koala habitat by Cherish the Environment Foundation Ltd and managed by agricultural scientist Mr Geoff Faulkner from Faulkner Consulting and foresters from Private Forestry Service Queensland (PFSQ). This large, ninety-hectare site included sixty-five hectares of new, five-year-old plantings of eucalyptus trees and old growth revegetation areas. The area is bounded by Warrill Creek to the east and cattle and hobby farms on its other borders. A seasonal creek and billabong also defined the site. The revegetation is under contract to Queensland Rail and the Department of Transport and Main Roads. The plantings of 34,000 thousand trees in 2015 consisted of Australian native trees that provide suitable habitat for koalas, the dominate plantings being *Eucalyptus tereticornis, Eucalyptus molluccana and Corymbia tessellaris* (see Table 1). The site was monitored for fauna movements and forms part of a wildlife corridor for koalas and other animals throughout the region.

**Table 1.** Numbers of each native Australian tree species planted at Mutdapilly, Ipswich, Queensland, to provide revegetated koala habitats.

| Species | Number planted at Mutdapilly |
| --- | --- |
| *Eucalyptus tereticornis* | 18,000 |
| *Eucalyptus moluccana* | 9,000 |
| *Eucalyptus crebra* | 2,000 |
| *Corymbia tessellaris* | 3,000 |
| *Eucalyptus melanophloia* | 500 |
| *Melaleuca irbyana* | 500 |
| *Corymbia intermedia* | 500 |
| *Melaleuca viminalis* | 300 |
| *Casuarina cunninghamiana* | 200 |
| Total | 34,000 |

The ten cameras were installed around the property, one metre above the ground for twenty-one weeks from Autumn to Spring. Sites were selected as those with potential to capture fauna movement, reduce the number of pictures taken due to flora blowing in the wind and to ensure good distribution of cameras across the site. Figure 1 shows an aerial shot of the Mutdapilly koala offset terrain and lists the GPS locations of each of the camera installation sites. Verbatim SD cards (32 GB) were replaced approximately every two weeks and the photographs were saved by camera number in Seagate 2 TB external hard drives.

**Figure 1.**
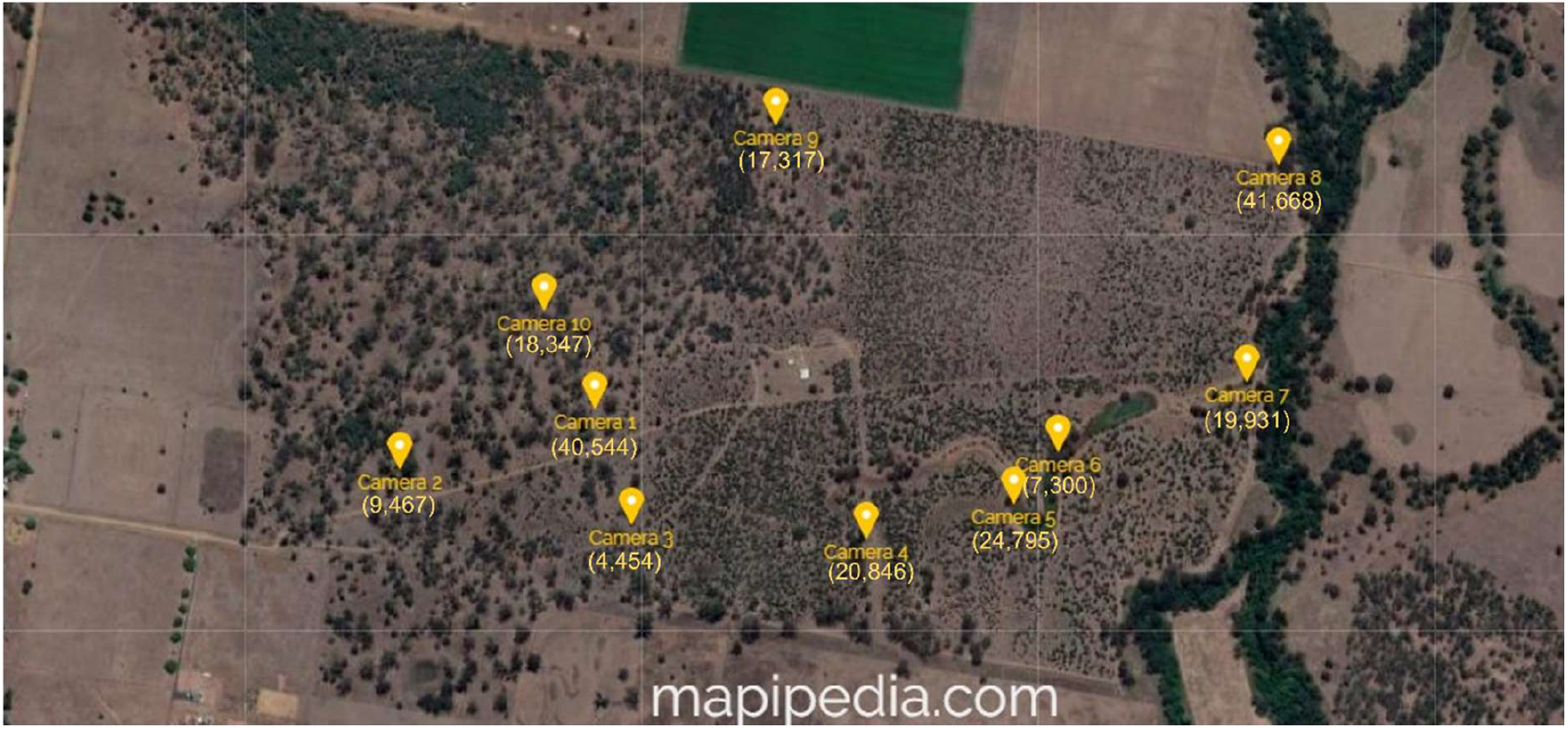
**Satellite image of the Mutdapilly koala offset site showing the locations of, and (in parenthesis) the number of photos captured by, the ten infrared motion sensor cameras. The image was created using mapipedia.com. Key landscape features such as the surrounding cattle properties, the creek and water course (under the densely grown tree line) are evident. GPS locations are as follows: camera number (latitude, longitude):** 1 (−2774892, 152.67637), 2 (−27.749647, 152.6736528), 3 (−27.75034, 152.67688), 4 (−27.7505, 152.68012), 5 (−27.75008, 152.68216), 6 (−27.74944, 152.68278), 7 (−27.74858, 152.68541), 8 (−27.74591, 152.68585), 9 (−27.74542, 152.67888), 10 (−27.7477, 152.67567).

During an excursion, twelve primary school students from Goodna State School investigated the heights and health of trees at each site, and made video recordings of features in the landscape. These students promoted the citizen science project with their peers at a school assembly for National Science Week and the entire school participated in ‘Koala Capture Week’, and the children, aged five to twelve years old, analysed the motion sensor photographs in a mini-citizen science project.

In order to analyse the presence or absence of animals within an image and conduct preliminary identification of the fauna shown, a website was created by partner and programmer Dr David Philpot using Python 2.7.16 (Version2.7.16:413a49145e, March 4 2019). A local computer was converted into a server to run the website to the school. Whenever an image was viewed and catalogued, a csv file was auto-populated with the data from the interaction with the website. Students at Goodna State School logged into the IP address of this server to access images. Each time a child selected “submit”, a new image was populated on the website. Some sample screen shots of the website are shown in Figure 2 with the initial question: “Can you see an animal in this picture?” If an animal is present, additional parameters appeared for students to complete: “What animal is it? Can you classify it?” with options such as koala, kangaroo, giant wood moth, bird (parrot, finches), possum, cow, feral pig, feral fox, lizard/reptile and other as illustrated in figure 3. A table for recording the temperature and atmospheric pressure from the image also appeared on screen. Throughout the project, regular communication between the coordinating teachers and offset site managers was maintained particularly with reference to feral animal presence. The analysis of the data generated via the use of the citizen science website is reported below.

**Figure 2.**
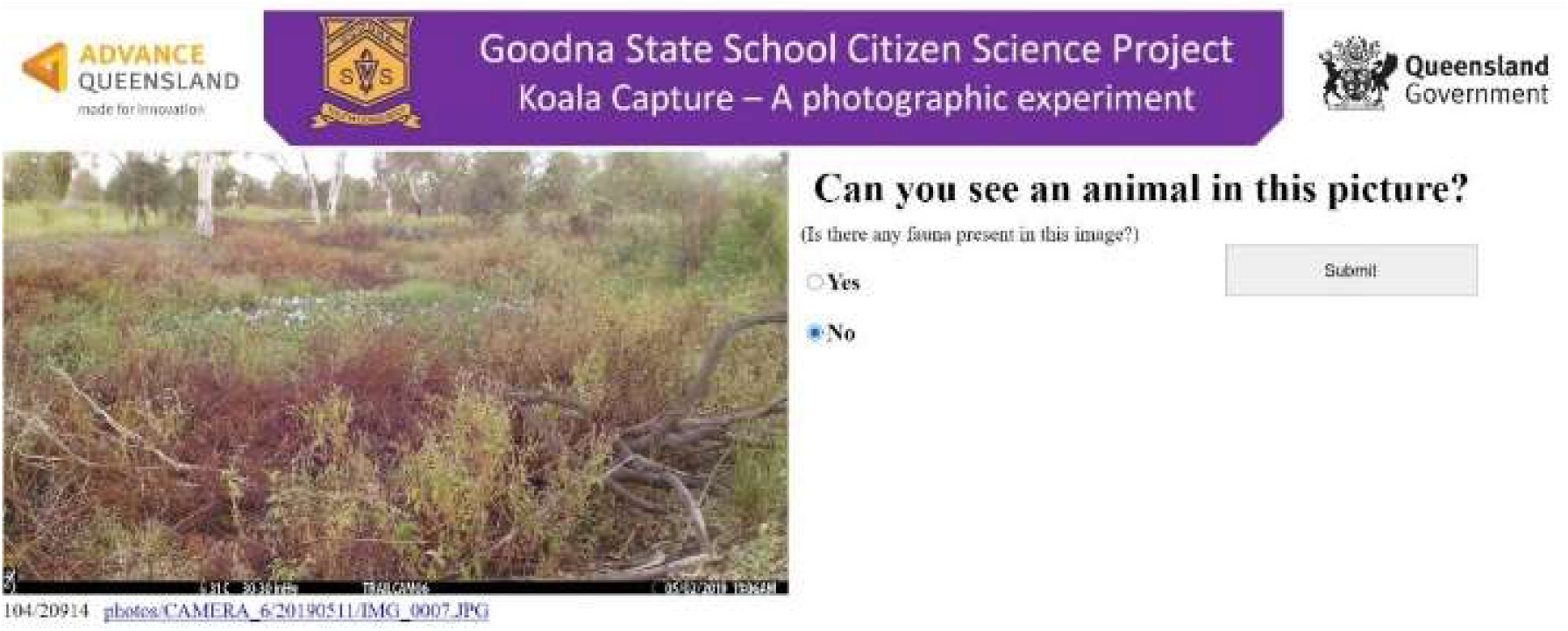
Sample screenshot of the ‘Koala Capture’ website showing the format when no animal is present in the image.

**Figure 3.**
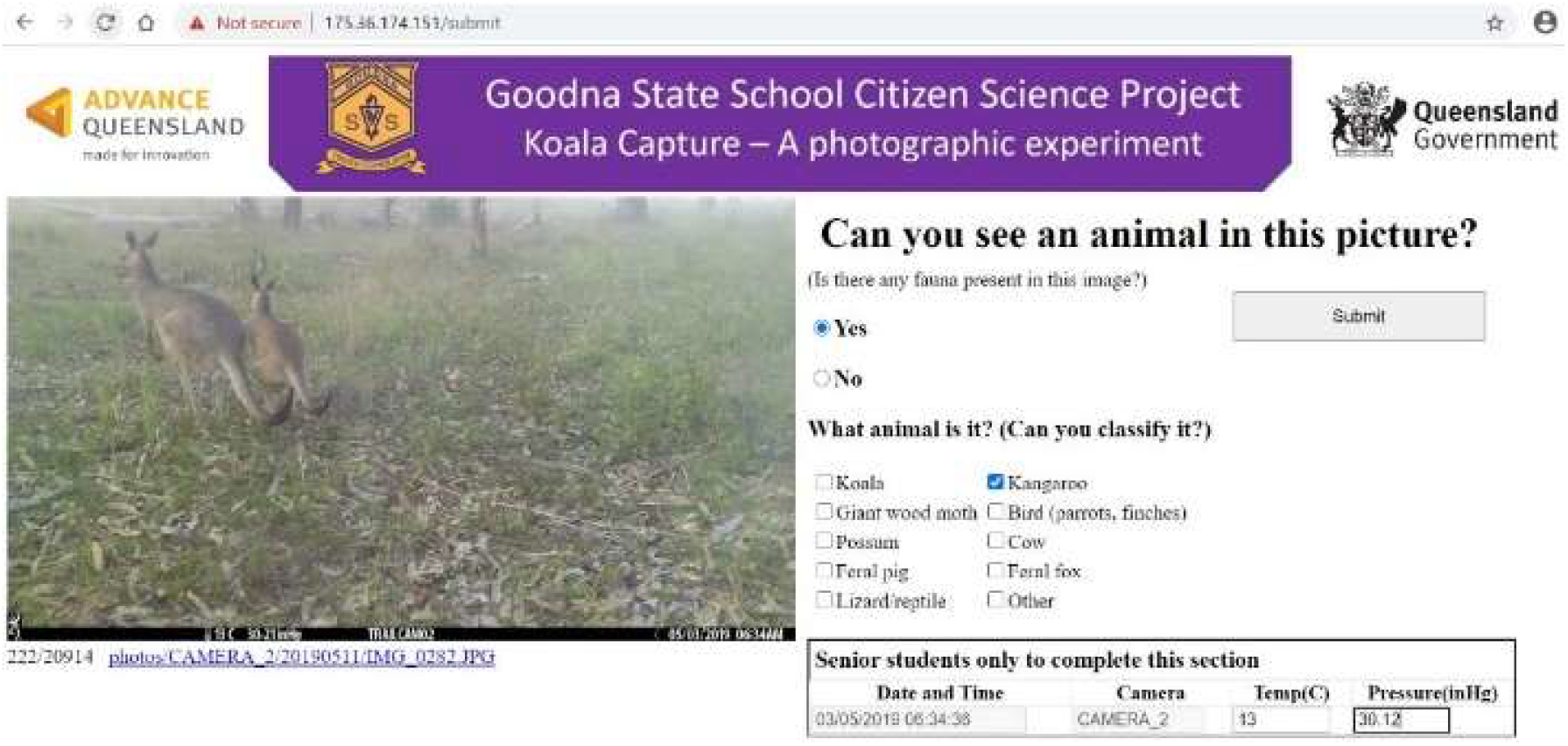
Sample screenshot of the ‘Koala Capture’ website showing the format when an organism is visible and additional detail that appears for student entry.

## Results

The ninety-hectare koala offset site at Mutdapilly in south-east Queensland had ten infrared cameras placed on trees throughout the landscape, taking in the water hole, neighbouring fence line with cattle grazing land, old growth areas and new plantings (Figure 1).

Some cameras took many photographs of plant movement agitated by the wind and may have filled the SD cards with numerous images within just a few days. Others, depending on the terrain, took less images within the same timeframe. This is common in ecological studies and modification of the camera settings and sensitivity to movement can assist in reducing these redundancies when looking for fauna.

### Goodna state school students – image analysis in ‘Koala Capture week’

More than 204,669 photographs were captured by the ten cameras over the twenty-one weeks. During ‘Koala Capture Week’ students searched images for the presence of fauna and attempted to identify them. Through the website, 13,132 photos (6.4% of total photos captured) were analysed by approximately 650 students in their weekly, one hour, science lesson. Students in the preparatory year (five-year-olds) accessed the website as a whole class, students in Grades 1-3 worked in pairs, whilst students in Grades 4-6 worked individually. Students took approximately five seconds to catalogue each photograph by interacting with the website to analyse the images provided. Each computer logged into the server’s IP address and an image was populated on the website. Students never saw the same image as their peers, however, due to the sequential nature of the images being populated on screen, their neighbour may have had the image immediately before, or after the photograph they saw. This was particularly useful if an animal moved quickly through a shot, or was difficult to identify (for example kangaroos hopping rapidly across screen, or a close up of a cow’s ear). The previous or subsequent photograph might show the kangaroo pausing to eat, or the whole face of a cow. In this way, working together, the students were more likely to correctly identify the animal present. However, given the age of the students, errors in identification were still inevitable.

Photographs taken by the cameras became infrared images at night and each image stored the temperature, atmospheric pressure, camera number, moon phase, date (dd/mm/yyyy) and time. An example image is in Figure 4 showing an adult koala transporting a baby across the landscape at 1:01am on the 10th of July 2019. The comma-separated values (csv) file generated by the website interaction automatically saved the image file name, camera number, internet protocol (IP) address of the computer that viewed the image, date and time of the image taken and the classification date. Once a student interacted with the image, the csv file also recorded the student’s entries – presence or absence of an animal, animal type, temperature and air pressure. This csv file was saved on the server computer.

**Figure 4.**
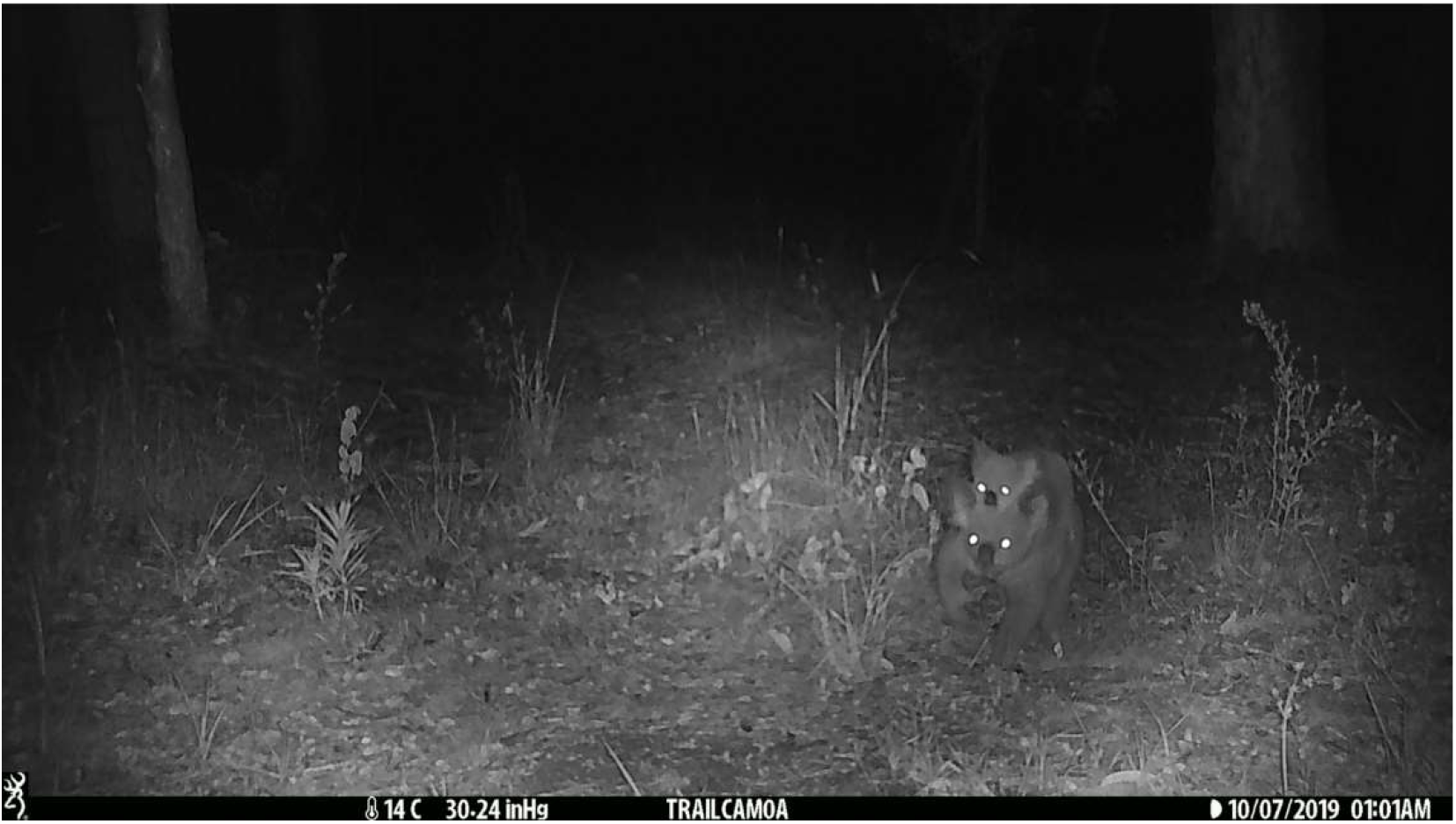
Example image from Camera 10 (X). This night time infrared image showing adult and juvenile koalas walking across the landscape, highlights the data present on each photograph: temperature, atmospheric pressure, moon phase, date and time.

The students who participated in this mini citizen science project ranged in age from four to twelve years old. The five day ‘Koala Capture Week’ resulted in the analysis of 13,132 images, with most students using the website for an hour’s session. Whilst the whole school (632 students) participated in the project, younger ages worked in groups of various sizes, thus reducing the number of computers needed overall for the investigation. However, on average, each primary school class analysed 412 images/hour. Over the six months that cameras were installed on-site, a total of 204,669 images were captured. At this rate of analysis, it would take a single class approximately 500 hours to analyse all the images. Or conversely, 500 classrooms could analyse all the images in just one hour (assuming the website could handle that many individuals accessing the server).

### Ecological results at the Mutdapilly offset site

This mini citizen science project engaged more than six hundred students in fauna identification at the Mutdapilly offset site via the use of infrared motion sensor cameras and the “Koala capture” website. As with all ecological projects with motion sensor cameras, an animal may be present in the view of the camera for a short time, and therefore, in a small number of images captured, or, in the case of the kangaroos, may linger around the camera in view: eating, lying down or walking slowly across the terrain. The images analysed by the students covered an approximately six-week period spanning May and June. Table 3 highlights the number of different dates each animal was sighted in the website by the children.

**Table 3.** Number of photographs of each animal type compared to the number of days the animal was captured during ‘Koala Capture Week’.

| Fauna | Number of images with animal present | Number of days animal was photographed | Average images/day |
| --- | --- | --- | --- |
| Koala | 28 | 12 | 2.3 |
| Kangaroo | 2,808 | 27 | 104.0 |
| Giant wood moth | 49 | 18 | 2.7 |
| Birds | 472 | 25 | 18.8 |
| Possum | 31 | 14 | 2.2 |
| Cow | 14 | 8 | 1.8 |
| Feral pig | 72 | 16 | 4.5 |
| Feral fox | 122 | 19 | 6.4 |
| Lizard/Reptile | 43 | 20 | 2.1 |
| Other (including humans) | 156 | 15 | 10.4 |

Some animals were photographed across the terrain, as illustrated by the number of times an animal was identified at a different camera location. For example, koalas were sighted more frequently at Cameras 1 and 10 but were identified in all other camera locations except for Camera areas 3 and 4. The sheer volume of kangaroo photographs, their rapid mobility and their ground dwelling nature, meant whilst they had regions they preferred to graze and travel through, the entire site captured kangaroo presence throughout the project. Camera 1 however was the most likely location to capture these marsupials.

The time-of-day data recorded by the camera and captured in the csv file also provides insight into the movement of animals around the site (table 4). Koalas were identified as being active across the day, however, errors in identification and potentially some ‘wishful-thinking’ is likely with students of this age. Feral fox and feral pig movements were interesting as both pest species were most active at night (between 7pm and 4am) but significant movement around the property also occurred around sunrise and throughout the day. Foxes appeared all across the site, but predominately near Cameras 7 and 1, whilst feral pigs also roamed widely with most common sightings near Camera 5 and 8 (data not shown). Understanding feral animal movements can aided in pest management at Mutdapilly.

**Table 4.** Number of sightings of various animals during various times of day. Raw data is listed first and expressed as a percentage of total images of that animal (in brackets).

| Time of day | Koala | Feral pig | Feral fox |
| --- | --- | --- | --- |
| Sunrise (4am-7am) | 3 (12.5%) | 12 (16.2%) | 22 (17.7%) |
| Daylight (7am-4pm) | 15 (62.5%) | 12 (16.2%) | 24 (19.3%) |
| Sunset (4pm-7pm) | 5 (20.1%) | 9 (12.1%) | 13 (10.4%) |
| Night (7pm-4am) | 1 (4.2%) | 41 (55.4%) | 65 (52.4%) |

### Student participation

Student interest in the project spanned all participating year levels from Prep to Year 6. Most children persevered through ‘boring’ photographs and cheered when an animal appeared in their photograph. The most enjoyable aspect for the citizen scientists was finding animals and looking at the different landscapes as reported to supervising teachers. However, errors in identification did occur. This is illustrated most stridently in the case of the Giant Wood Moth (*Endoxyla cinereus*), an insect whose larvae eat Eucalyptus wood. Figure 5 shows a blurred image of the nocturnal moth flying past a camera at 1:05am. The wingspan of the moth can be up to twenty-three centimetres across and the flapping rate would be faster than the shutter speed at night resulting in blurred images. This image was identified by one of the teachers after scanning through camera pictures. The teacher’s experience with this insect allowed for confirmation of the identity of the animal. However, when cataloguing the photographs through the website, approximately half of the images classified as “giant wood moth” by students, occurred in photographs taken during the day. The giant wood moth is nocturnal – highlighting the importance of an experienced eye being cast over the positive images (data not shown).

**Figure 5.**
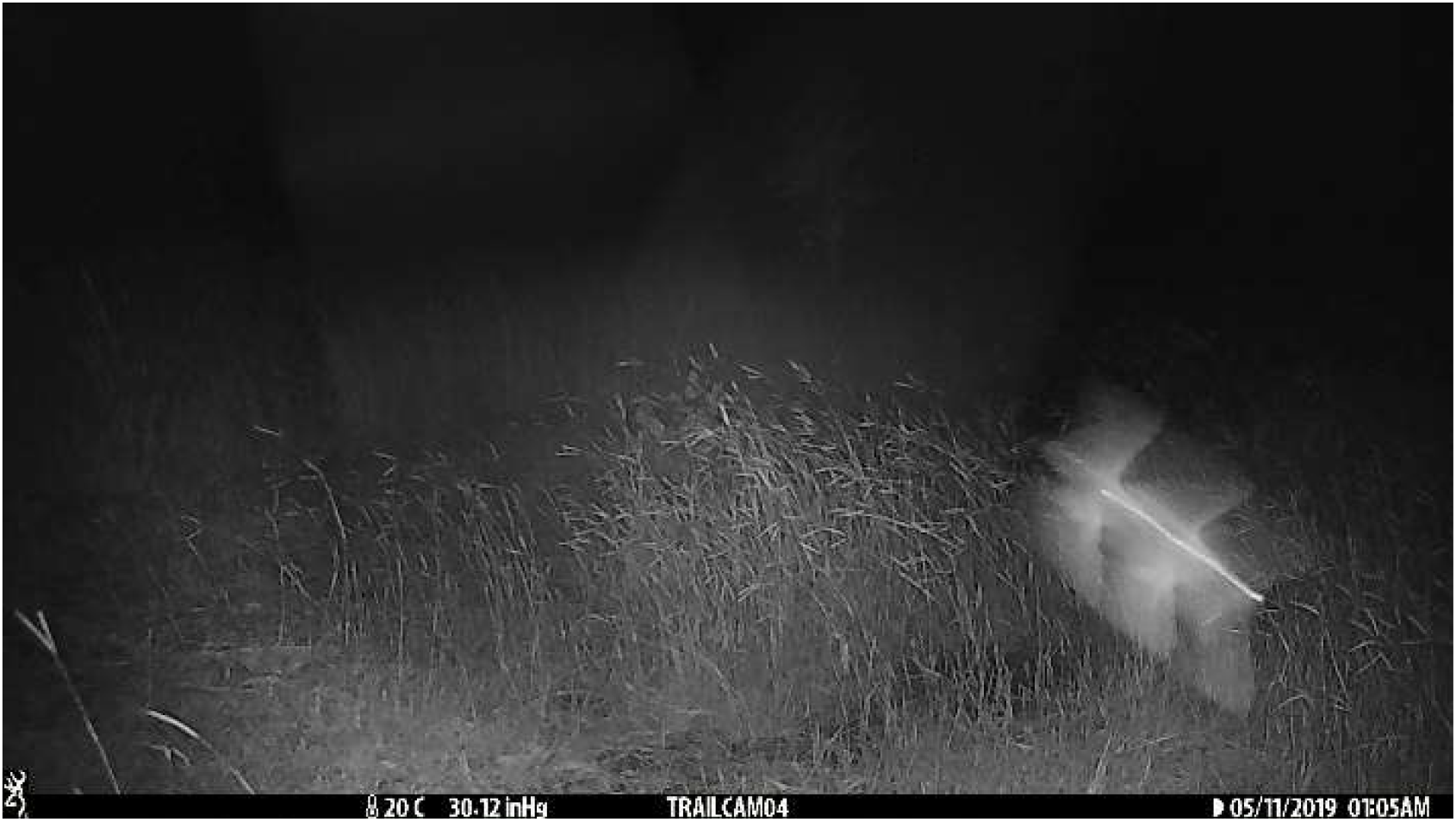
A giant wood moth (*Endoxyla cinereus*) flying past Camera 4 at 1:05 am, 11 May 2019. This nocturnal insect spends most of its lifecycle inside the stems of host eucalypts. The thirty-gram adult moth only emerges at night to lay its 20,000 eggs. Some students erroneously identified moth presence during the daytime images.

## Discussion

Habitat destruction, bushfires, vehicular traffic and diseases have decimated koala populations in the eastern states of Australia. Consequently, koala revegetation sites and wildlife corridors are fundamental for the marsupial’s survival. This mini project showed it was possible to use primary school children as citizen scientists to detect koala (and other fauna) presence using web-based systems as long as the system was simple to use. However, some limitations and constraints to the project must also be addressed.

### Successes: ecological management, student participation and image analysis

Koala habitat revegetation programs are designed to facilitate koala population growth by their placement in wildlife corridors, the tree species planted and the ongoing management of these environments. The Queensland government legislation outlined in the Queensland Environmental Offsets Policy (Department of Environment and Science 2021) mandates the creation and regulation of offset sites for koala habitats. Ecological management of these sites by experts (including Faulkner Consulting and Private Forestry Service Queensland) enhances sustainable practice for koala conservation as they bring specialist knowledge to the planting and maintaining of plant species in degraded or arid soils (Private Forestry Service Queensland 2020). Promisingly, the five-year-old plantings at the Mutdapilly koala offset site studied here have shown evidence of koala movement into the area through visual sightings, scat presence and characteristic tree scratchings in what was previously arid grazing land that had been cleared for beef cattle production (personal communication Faulkner Consulting 2021). This mini citizen science project informed property management of ecology at the Mutdapilly koala offset site. When the teachers involved in this project uploaded photos onto the database for the website, any feral animal presence observed was shared immediately with Faulkner Consulting. For example, cameras captured feral pig movements along the fence line of the property enabling site managers better understand the transient nature of pig incursions into the property. Council wildlife officers determined that pig access was transitory and they roamed up and down the creek causing damage to the watercourse banks. Similarly, feral foxes were captured by the motion sensor cameras and seemed to confirm that earlier control measures had an ongoing impact. The onsite tenant also reported sightings of feral foxes but predation impact on koalas did not appear to be widespread. Baiting was not implemented due to the proximity of neighbours and domestic animals (personal communication Faulkner Consulting 2021). Students at Goodna State School loved hearing stories about feral animal presence and their impact on the Mutdapilly site (as shared by science teachers with their classrooms).

The student citizen scientists were highly excited to participate in the ‘Koala Capture Week’. Students appeared focussed during the image analysis sessions. Students often expressed exclamations of joy at animal presence and teacher discussions with students after ‘Koala Capture Week’ indicated most students thoroughly enjoyed the experience. The use of primary/elementary school students as citizen scientists, whilst not unique, is not commonplace. Some citizen science projects scout for participants through advertising special events (e.g. annual FrogID week, FrogID, 2020), or by adding projects to specialised citizen science association project finders (e.g. The Australian Citizen Science Association (ACSA) project finder managed by Atlas of Living Australia, 2021) some of which are simple enough for young students to use. The project reported here is the first time in south-east Queensland that a partnership has used motion sensor cameras to gather information on koala (and other fauna) presence utilising the engagement of a local primary school as the citizen scientists. Another local and contemporary research-citizen-science-school-partnership includes the analysis of native grass pollen data in south-east Queensland (Van Haeften et al. 2021). Year nine agricultural science students at a Brisbane secondary school used an online app to collect and submit native grass distribution information around their school and homes during the covid-19 lockdowns of 2020. This facilitated a two-way knowledge exchange between the secondary and tertiary education sectors (Van Haeften et al. 2021) and fostered interest in research. In our project, the young children were highly motivated and loved looking for animals on the ‘Koala Capture’ website, assisted by the iconic and beloved status koalas hold in Australian’s hearts and minds (Jackson 2010).

Whilst citizen science projects provide benefit to the landholder, scientist or research organisation by dividing up time consuming, repetitive tasks amongst large numbers of participants, the educational benefit to the participant cannot be understated. According to Bonney et al. (2009) studies show benefits to participants from participation in citizen science projects, including, but not limited to: increased awareness and/or understanding of key scientific concepts, knowledge gains of the process of science and increased science inquiry skills (Bonney et al. 2009). Participation in research through citizen science projects creates authentic learning experiences, fosters inquiry, develops place-based nature experiences and elevates the public’s understanding of science, the environment, and stewardship of the Earth (Dickinson and Bonney 2012; Shirk *et al*. 2012). It is hoped that including primary school students as citizen scientists in projects such as this ‘Koala Capture’ project will foster a broader appreciation for scientific research and a love for the animals and plants with which humans share this planet.

The fauna identified by students through the ‘Koala Capture Week’ included kangaroos, koalas, possums, birds, moths, cattle and reptiles as well as feral pigs and foxes. The analysis of photographs generated by motion sensor cameras in ecological studies can be a time-consuming task and the use of citizen scientists to perform preliminary scans of photos has merit if appropriate checking measures are implemented by the research head. An exciting aspect of the investigation of the fauna present in a koala revegetation offset site at Mutdapilly by Goodna State School students meant the children contribute to environmental management from their local area (i.e. Mutdapilly is thirty-eight kilometres from Goodna and less than thirty minutes’ drive away). This local agency was motivating for students involved particularly those who had physically attend the site and mounted cameras. Technologically, whilst many high schools have the coding capability to generate their own citizen science websites, Goodna State School was indebted to the expertise and generosity of David Philpot for creating the database infrastructure and supporting the school science teachers to manage the web site, maintain the server on a private computer and troubleshoot any connection issues. This mini project showed that young students could engage in citizen science ecological research, provided the technological interfaces were simple and the teachers could easily implement the program into the school setting.

### Challenges: technological limitations, student error and time constraints

Whilst this mini citizen science project utilised the time and identification skills of primary school students in a productive manner, it was not without its challenges. Some citizen science projects benefit from the expertise of “super citizen scientists” as described by Hames et al. (2002) who obtained two hundred highly engaged citizen scientists by recruiting from the Birds in Forested Landscapes participant base (Hames et al. 2002). However, the project reported here relied on the engagement and expertise of four to twelve-year-old students enrolled in the local primary school. Students of such a young age inevitably display observer variability. Teachers reported that some students glanced at each image, rapidly deciding if any fauna was present, whilst others laboriously zoomed in on each image and scanned the grasses diligently. Other students were so enthusiastic to identify a snake in the grass, that many a log, branch or fallen twig was erroneously declared a reptile. It has already been noted that some of the koala ‘sightings’ were a results of student wishful thinking, and giant wood moth sightings during the day are unlikely in this noctural animal. However, such biases are not new, nor unique to citizen science projects with young participants. According to Weir et al. (2005), analysis-related challenges in citizen science projects such as sampling bias, observer variability, and detection probability are not easily addressed with statistical hypothesis testing or model selection approaches (Weir et al. 2005). Philippoff and Baumgartner (2016) analysed the effect of inconsistent, erroneous and unusable data in an intertidal citizen science project in Hawaii. They grouped errors into four broad categories of missing data, sloppiness, methodological errors, and misidentification errors and reported decreased error rates with field trip experience and student age (Philippoff and Baugmgartner 2016). Training in identification, education on the importance of accurate data and investment in the research outcome helped reduce error rates.

The ten motion sensor cameras at the Mutdapilly koala offset site collected a large amount of data. The cameras remained on location for twenty-one weeks resulting in 204,669 images. However, during ‘Koala Capture Week’, only six weeks’ worth of images were studied by the primary school students, with fifteen weeks of images remaining un-investigated. Preliminary scans of the images on SD cards revealed whole days where wind movement of the plants surrounding the cameras filled the memory card, and the decision was made by the teacher authors to cull a percentage of these ‘empty’ photographs so as to decrease potential student boredom and maintain on task behaviour during the analysis lessons. This action skewed the data reported here with reference to photographs analysed to real numbers of fauna present but was necessary given the time constraints on the project. However, it must be noted that no images were actually deleted from the research database. Larger scale versions of this project would benefit from running the server through Education Queensland’s infrastructure. Whilst this would require more time for vetting and setting up of the IT systems, it would allow multiple public schools to be involved in the project, more simultaneous users to access the website without causing large delays loading images, and the analysis of more images without the need for this artificial vetting.

The website created for ‘Koala Capture Week’ was easy to use for even the smallest of fingers and youngest scientists. The server-web site system reported here worked effectively for approximately sixty simultaneous users without significant delays loading images. However, there are safety considerations when opening such projects up to broader IT systems and other student users. In Queensland, The Information Privacy Act (2009) legislates how the Department will collect, store, use and disclose personal information about students (Queensland Government 2019). For example, in pollen project reported by Van Haeften et al. (2021) students had to provide their personal details to download the citizen science app and Education Queensland had to both approve this process and be informed of the country in which the data was stored, to ensure the security of student personal information. At the local level, school approvals were required by the information technology (IT) manager and Deputy Principal (Van Haeften 2021). In order to avoid these issues of personal student data being stored on overseas servers, this mini citizen science project did not require Goodna State School participants to create a profile to access the website. The only identifying information collected by the local server was the IP address of each user. The primary school students did use their personalised Education Queensland logins to use the laptops and access the internet, but given that the server running the website was simply another local laptop and turned off once the project was completed, the online risk was reduced. Approvals were sort to run the ‘Koala capture’ website in this way from Education Queensland IT managers and school specific permissions granted by the Principal of the school. If the project was expanded to include other schools and thereby analyse more photos, running the website on the Department of Education’s servers would be required and a risk analysis to student privacy performed, but this was beyond the scope of this project.

## Conclusion

The beloved nature of the koala, the close proximity of the Mutdapilly offset site and the usability of the ‘Koala Capture’ website combined to create a successful citizen science project using young students. The motion sensor cameras captured thousands of photos and students were able to scan images for fauna presence. However, it is essential that teacher’s provide education to student citizen scientists on the importance of accurate collection of data for a research utility. Engagement with larger numbers of primary school students would have allowed more images to be catalogued but future projects would require additional investment in server infrastructure and online student safety.

## Supplemental Files List

None.

## Data Availability Statement

The raw data and photos collected in this project has not been made publicly available as it remains the property of the researchers but may be requested by emailing the first author.

## Acknowledgments

Teachers from Goodna State School wish to acknowledge that this project would not have been possible without the support of School management staff, particularly Acting Principal Daniel Dempsey. Access to the Mutdapilly Koala Offset site by students facilitated by the generosity of owners, Cherish the Environment Foundation Ltd and kindness of agricultural scientist Mr Geoff Faulkner from Faulkner Consulting and foresters from Private Forestry Service Queensland (PFSQ) to allow the camera installation, student excursions and data collection on site.

## Funding

This project was made possible by an Advance Queensland Engaging Science grant of $10,000 (2018-2019) Mutdapilly Koala Offset, School Based Citizen Scientist Project Goodna State Primary School. The grant facilitated the purchase of ten motion sensor cameras and covered project costs. The project aimed to foster environmental awareness in a primary school community that has emerging understandings of sustainability and to promote STEM awareness in the broader Queensland community. The Mutdapilly offset is a unique project and the school’s STEM program has handwritten endorsements from environmentalists Sir David Attenborough and Dr Bob Brown, and the CEO of the Australian Conservation Foundation.

## Authors’ Contributions Statement

**Sharon Williams**: Conceptualization (equal); Project administration (equal); Writing original draft (lead): Writing-review & editing (equal), Methodology (equal). **Gerard Salmon**: Conceptualization (equal); Project administration (equal); Writing original draft (supporting): Writing-review & editing (equal), Methodology (equal). **Daniel Dempsey**: Project administration (supporting); Writing-review & editing (supporting). **David Philpot**: Conceptualization (supporting); Writing original draft (supporting): Writing-review & editing (equal), Methodology (supporting). **Geoff Faulkner**: Conceptualization (equal); Project administration (supporting); Writing original draft (supporting): Writing-review & editing (equal), Methodology (supporting)

## Competing Interest Statement

There are no competing interests for this project.

## Ethics Clearance

The data reported in this paper was gathered as part of normal teaching and learning practices at the Queensland Department of Education school, Goodna State School, and as no identifying information, individualised student work, or personal information was collected or reported, no human ethics clearances were required.

## Notes

### Competing Interest Statement

The authors have declared no competing interest.

